# IgA/IgM chromatographic depletion enables efficient 20-nm virus nanofiltration of mini-pool caprylic-acid IgG

**DOI:** 10.64898/2026.02.26.708374

**Authors:** Liling Delila, Daniel Strauss, Thierry Burnouf

**Affiliations:** Graduate Institute of Biomedical Materials and Tissue Engineering, College of Biomedical Engineering, Taipei Medical University, Shuang-Ho campus, New Taipei City, Taiwan; Asahi Kasei Bioprocess America, Inc., Glenview, Illinois, USA; International PhD Program in Biomedical Engineering, College of Biomedical Engineering, Taipei Medical University, Shuang-Ho campus, New Taipei City, Taiwan

**Keywords:** Virus safety, Caprylic acid precipitation, Nanofiltration, Resource-limited settings

## Abstract

Global shortages of human plasma-derived immunoglobulin G (IgG) remain a major challenge for treating primary immunodeficiencies, especially in low- and middle-income countries. Ensuring virus safety is essential, and nanofiltration provides robust removal of small, non-enveloped viruses. We examined whether removing immunoglobulin A (IgA) and immunoglobulin M (IgM) by anion-exchange chromatography improves the performance of 20-nm nanofiltration applied to small-pool caprylic acid–purified IgG. Cryo-poor plasma was treated with 5% caprylic acid at pH 5.5, concentrated by ultrafiltration, and processed on Fractogel TMAE to deplete IgA and IgM. The IgG flow-through was filtered sequentially through Planova 35N and 20N (or S20N) filters. Direct nanofiltration of caprylic acid–treated IgG with residual IgA and IgM led to rapid membrane clogging and low throughput. Depletion of IgA and IgM increased filtration capacity more than threefold and stabilized flux. Dynamic light scattering confirmed the predominance of monomeric IgG and absence of aggregates after chromatography and nanofiltration. Overall, this process combines two complementary virus reduction steps, caprylic acid treatment and nanofiltration, and provides a practical option for LMICs to convert available domestic plasma into IgG; it could also be adapted to the manufacture of hyperimmune or convalescent IgG preparations.

## Introduction

Access to plasma-derived medicinal products (PDMP), including immunoglobulin G (IgG), remains highly unequal worldwide (Belmonte et al., 2025; Prevot and Jolles, 2020). This reflects limited plasma collection and fractionation capacity, rising use of IgG for immunomodulatory indications in high-income countries (HICs), and economical and yield constraints of current large-scale manufacturing technologies (Burnouf, 2007; Curling et al., 2025; Radosevich and Burnouf, 2010). In addition, in many low- and middle-income countries (LMICs), recovered plasma produced from whole blood is underused or discarded because local processing capacity is lacking, and contract or domestic fractionation is often not feasible when plasma volumes and quality systems are insufficient (WHO, 2021).

For patients with primary and secondary immunodeficiencies, IgG replacement is lifesaving. Yet, despite growing global demand, many LMICs face persistent shortages of this essential medicine (El Ekiaby et al., 2024). Most IgG is produced and distributed in HICs, leaving LMICs with limited access because of cost, limited supply, and constrained infrastructure.

Recent World Health Organization (WHO) guidance and statements from major transfusion organizations emphasize that improving access to safe plasma protein products should be a global health priority (Burnouf et al., 2024b; Burnouf et al., 2022; WHO, 2021). Beyond strengthening plasma collection and quality systems, there is a need for technologies that allow stepwise development of local processing capacity. In this context, small-scale and modular approaches to virus-reduced plasma-derived medicinal products (PDMPs), including immunoglobulins (El-Ekiaby et al., 2015) and mini-pool plasma/cryoprecipitate processing (El-Ekiaby et al., 2010), can provide pragmatic interim options for access to treatment when industrial fractionation or contract manufacturing is not yet possible (Burnouf et al., 2024b; Burnouf et al., 2020; WHO, 2021).

A practical example is the caprylic acid (CA) precipitation process developed in Egypt to isolate immunoglobulins from cryo-poor plasma (El-Ekiaby et al., 2015). The process is technically straightforward, can be implemented in closed bag systems with basic equipment, and the CA step provides effective inactivation of lipid-enveloped viruses. Clinical studies reported pharmacokinetics and efficacy similar to commercial IgG in immune thrombocytopenia (Elalfy et al., 2017) and in children with primary immunodeficiency (Selim et al., 2025). A larger-scale adaptation has also been used to produce anti–SARS-CoV-2 immunoglobulins in Pakistan (Ali et al., 2021a), with clinical evaluations reported (Ali et al., 2022; Ali et al., 2021b). More broadly, a similar CA fractionation process is used to manufacture antivenom immunoglobulins from hyperimmunized horse plasma (Rojas et al., 1994), which are listed as essential medicines for adults and children by the WHO (WHO, 2023).

However, CA-based IgG purification has some limitations (Burnouf et al., 2024a; Pergent et al., 2024). CA treatment does not efficiently inactivate or remove non-enveloped viruses, so an additional orthogonal virus reduction step is needed, particularly when scaling to larger pools which have more risks to include undetected contaminated plasma donations (Burnouf and Radosevich, 2000). In addition, CA IgG intermediates still contain substantial IgA and IgM (Ali et al., 2021a; El-Ekiaby et al., 2015). These high-molecular-weight immunoglobulins may contribute to adverse reactions in some patient subgroups (Radosevich and Burnouf, 2010). Furthermore large molecules like IgM can hinder downstream virus removal by small-pore nanofiltration, a processing step that is especially effective for the removal of blood-borne small non-enveloped viruses such as parvovirus B19 and hepatitis A virus (Burnouf and Radosevich, 2003; Roth et al., 2020a), by accelerating filter fouling and reducing capacity and throughput. This constrains implementation of robust dual virus-reduction strategies (Burnouf, 2007; Velthove et al., 2013).

We hypothesized that anion-exchange chromatography to selectively deplete IgA and IgM from CA-purified IgG (Cheng et al., 2021; Wu et al., 2013) would improve small-pore nanofiltration performance and thereby strengthen virus safety while supporting stepwise scale-up. This proof-of-concept laboratory scale-down study evaluated the impact of IgA/IgM removal by anion-exchange chromatography on sequential nanofiltration in a small-pool CA-based IgG process, as a potential stepwise pathway to better utilize domestic plasma resources in LMICs (WHO, 2021).

## Materials and Methods

### Plasma Collection and Preparation of Cryo-poor Plasma

A pooled preparation of human plasma units (up to a total volume of 2 L), anticoagulated with citrate were obtained from regular whole-blood or apheresis donors under Institutional Review Board of Taipei Medical University (TMU-JIRB N201802052) approval as previously described (Cheng et al., 2021). Plasma was separated, frozen within 8 h of collection, and stored at ≤ –20 °C until use. For processing, units were thawed at 2–4 °C for ∼16 h. Cryoprecipitate was removed by centrifugation at 10,000 × g for 30 min at 2–4 °C using Heraeus Multifuge X1R centrifuge, (Thermo Fisher, MA, USA). The supernatant (“cryo-poor” plasma; CPP) was collected for further fractionation.

### Caprylic Acid (CA) Precipitation

CA fractionation was performed as previously described (Cheng et al., 2021; El-Ekiaby et al., 2015). Briefly, CA (Merck, KGaA, Darmstadt, Germany) was added slowly to CPP under vigorous stirring to a final concentration of 5% (v/v). During CA addition, the pH was adjusted and maintained at 5.5 ± 0.2. The mixture was incubated at 22±2 °C for 90 min with mild agitation. Precipitated proteins were removed by centrifugation at 10,000 × g for 30 min at room temperature. The supernatant containing immunoglobulins (CA-IgG) was clarified by sequential filtration through 1.2 µm and 0.22 µm filters (Milligard PES and Durapore 0.22 µm; Merck) (Cheng et al., 2021).

### Buffer Exchange and Sample Preparation

CA-IgG was buffer-exchanged into 25 mM sodium acetate, pH 6.0 using tangential flow filtration (TFF) (Pellicon XL cassette, 30 kDa MWCO; Merck Millipore). 2L of pooled plasma were used for CA precipitation, yielding 1.7–1.8 L of CA-IgG, and subsequently concentrated by approximately 50% using TFF. The concentrate was subsequently clarified using a depth filter (Millistak+ HC pod; Merck) followed by a 0.22 µm PES filter prior to chromatography.

### Removal of IgA and IgM by Fractogel TMAE Anion-Exchange Chromatography

Anion-exchange chromatography was performed using Fractogel® EMD TMAE (M) resin (Merck KGaA, Darmstadt, Germany). A 5 mL Fractogel TMAE (M) prepacked column (MiniChrom format; Merck), was equilibrated with 25 mM sodium acetate buffer (pH 5.9). EQ-IgG, corresponding to CA-IgG previously buffer-exchanged into 25 mM sodium acetate, pH 6.0, was loaded at a linear flow rate of 180 cm/h. A total of 62.5 mL of EQ-IgG with IgG concentration ∼8.0 mg/mL was applied, corresponding to an IgG load of approximately 100 mg IgG per mL of resin. After sample loading, columns were washed with equilibration buffer. The flow-through fraction containing IgG-enriched and depleted of IgA/IgM, was collected continuously and pooled based on the UV absorbance at 280 nm. The pooled flow-through was subsequently used for nanofiltration. After the flow-through collection, the column was washed with equilibration buffer and then subjected to a high-ionic-strength elution step using 500 mM sodium acetate (pH 4.5) to remove bound IgA and IgM and followed by a regeneration step with 1.5 M NaCl in 250 mM sodium acetate and cleaning-in-place (CIP) with 0.5 M NaOH (Cheng et al., 2021). After CIP, the column was equilibrated in a storage buffer and stored at 4 °C for further use.

### Nanofiltration

IgA/IgM-depleted IgG (Fractogel flow-through) was subjected to sequential virus nanofiltration using Planova 35N followed by Planova 20N or S20N filters (Asahi Kasei Bioprocess), each with an effective membrane surface area of 0.001 m². Prior to filtration, each nanofilter was pre-flushed according to the manufacturer’s instructions with the 25 mM sodium acetate buffer to remove preservatives and to equilibrate the membrane. The Planova 20N and S20N filtrations were performed at room temperature under constant transmembrane pressure at 0.8 and 2.20 kgf/cm², respectively, monitored and controlled using an ÄKTA Prime Plus system (Cytiva, Marlborough, MA, USA). Filtrate volume was recorded over time to derive filtration performance curves. Filtration flux was calculated as the filtrate volume normalized to membrane surface area and filtration time and is expressed in units of L·m⁻²·h⁻¹.

### Protein Quantification and Purity Assessment

IgG, IgA, and IgM concentrations in plasma and process intermediates were determined by immunoturbidimetry on a Cobas C502 analyzer (Roche Diagnostics, Mannheim, Germany). IgG integrity and purity were assessed by Sodium dodecylsulfate polyacrylamide gel electrophoresis (SDS-PAGE) under non-reducing conditions. 10 µg protein per lane was mixed with 4× sample buffer and heated at 70 °C for 10 min. Samples were separated on 4%-12% Bis-Tris gels (NuPAGE, Novex Life Technologies, CA, USA) at 70 V for 30 min followed by 200 V for 45 min. Proteins were stained using a Coomassie-based stain (InstantBlue, Abcam) and visualized by digital photography using a white background under uniform ambient lighting.

### Particle Size Analysis

The apparent size distribution and subvisible particulate content of IgG preparations (before and after nanofiltration) were assessed by dynamic light scattering (DLS) and nanoparticle tracking analysis (NTA). For DLS, samples were measured using a Zetasizer Nano instrument (Malvern Instruments) at 25 °C after 120 s equilibration in disposable polystyrene cuvettes. Measurements were performed in triplicate with ≥3 consecutive runs per sample. Scattered light was detected at 173°. Data are reported as Z-average hydrodynamic diameter and polydispersity index (PdI). For NTA, samples were analyzed using a NanoSight instrument (Malvern Instruments) to quantify the concentration and apparent size distribution of subvisible, light-scattering particles/aggregates in the IgG solutions. Samples were diluted in 0.22 µm-filtered ddH₂O to achieve a particle concentration within the instrument’s recommended range. For each sample, three 60-s videos were recorded under identical acquisition and analysis settings and processed using the manufacturer’s software. NTA results are reported as mean and mode diameters, and particle concentrations were calculated from tracked counts and dilution factors.

### Statistical Analysis

Data are presented as mean ± standard deviation (SD). Results were limited to descriptive statistics, including calculation of means, standard deviations, percentages, and graphical presentation of filtration performance curves. No formal hypothesis testing or inferential statistical analyses were performed, as the study was designed to evaluate process performance and feasibility rather than to test predefined statistical differences. The number of experimental replicates is indicated in the results section and figure legends.

## Results

### CA fractionation produces an IgG-rich intermediate that still contains IgA and IgM

CA precipitation of CPP generated an IgG-rich intermediate (CA-IgG) that still contained substantial levels of IgA (1.6 ± 0.2 mg/mL) and IgM (0.5 ± 0.1 mg/mL), with IgG at 8.7 ± 0.3 mg/mL. In unfractionated CPP, IgA, IgM, and IgG concentrations were 2.0 ± 0.1, 0.8 ± 0.1, and 10.2 ± 0.7 mg/mL, respectively. These results confirmed that CA precipitation enriches IgG but does not effectively deplete IgA or IgM, which remain at appreciable levels in the CA-IgG intermediate.

### Removal of IgA and IgM by anion-exchange chromatography

When the diafiltered CA-IgG fraction (EQ-IgG) was loaded onto a 5 mL Fractogel TMAE anion-exchange column equilibrated in 25 mM sodium acetate (pH 6.0), IgA and IgM were substantially depleted in the IgG flow-through fraction. IgA decreased from 1.11 mg/mL to <0.24 mg/mL and IgM decreased from 0.35 mg/mL to <0.20 mg/mL, while IgG in the flow-through remained >5.8 mg/mL, corresponding to an overall IgG recovery of 94.84% (Table 1). After nanofiltration, IgG concentrations were 5.27 mg/mL in the Planova 35N filtrate and 5.04 mg/mL in the sequential Planova 35N→20N filtrate, corresponding to overall IgG recoveries of 84.17% and 78.05%, respectively, while IgA and IgM remained below the limits of quantification (Table 1). A representative breakthrough chromatogram is shown in Figure 1: the UV280 trace shows the IgG flow-through during sample loading, whereas retained proteins (including IgA/IgM) were recovered during the high-salt elution/regeneration step.

**Figure 1.**
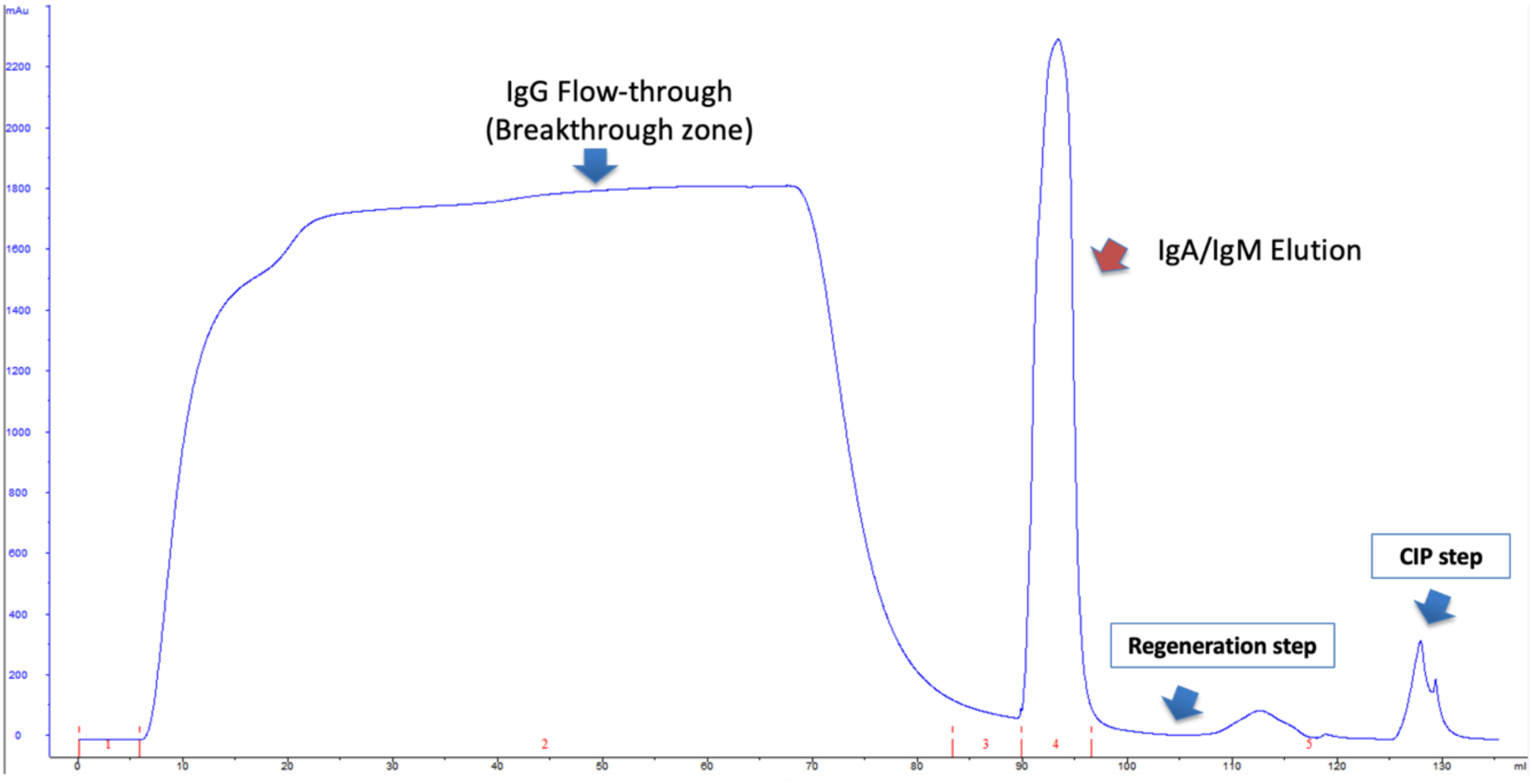
UV280 breakthrough chromatogram during anion-exchange chromatography on Fractogel TMAE (M) used to deplete IgA and IgM from the caprylic acid–purified EQ-IgG intermediate. The IgG-rich flow-through (breakthrough zone) was collected, whereas retained proteins (including IgA/IgM) were subsequently eluted during the regeneration step. The final peak corresponds to the column cleaning-in-place (CIP) step. UV absorbance is shown at 280 nm as a function of elution volume (mL).

**Table 1.** IgG, IgA, and IgM concentrations and IgG recoveries in the starting material (EQ-IgG), the Fractogel TMAE (M) flow-through, and after sequential Planova 35N and 20N nanofiltration. IgG content (mg) was calculated as concentration x volume. Step recovery (%) was calculated relative to the preceding step, and overall recovery (%) relative to the starting material. IgA and IgM values reported as “<” indicate concentrations below the assay’s limit of quantification.

| **step** | **IgG** | **IgA** | **IgM** | **Volume**<br>**(ml)** | **IgG**<br>**content**<br>**(mg)** | **Step**<br>**recovery**<br>**versus**<br>**previous**<br>**(%)** | **Overall**<br>**recovery**<br>**rate (%)** |
| --- | --- | --- | --- | --- | --- | --- | --- |
| **EQ-IgG**<br>**(starting**<br>**material)** | 8.02 | 1.11 | 0.35 | 50.00 | 401.00 |  |  |
| **Fractogel**<br>**TMAE (M)**<br>**Flow-through** | 5.83 | L<0.24 | L<0.20 | 65.23 | 380.30 | 94.84 | 94.84 |
| **Planova 35N**<br>**Filtrate** | 5.27 | L<0.24 | L<0.20 | 64.04 | 337.50 | 88.75 | 84.17 |
| **Planova 20N**<br>**filtrate** | 5.04 | L<0.24 | L<0.20 | 62.30 | 313.00 | 92.74 | 78.05 |

### Nanofiltration: filtration performance is improved by IgA/IgM depletion

Direct nanofiltration of EQ-IgG (containing high levels of IgA and IgM) through Planova 20N (0.001 m²) resulted in a rapid loss of filterability. As shown in Figure 2A, permeate flux declined sharply after ∼29 min, with only ∼16 mL of filtrate collected within 2 h at a transmembrane pressure of 0.8 kgf·cm⁻², corresponding to an average permeate flux of ∼8 L·m⁻²·h⁻¹ (Table 2). Interestingly, pre-filtration using Planova 35N before Planova 20N did not improve performance. Sequential filtration through Planova 35N (0.001 m²) followed by Planova 20N (0.001 m²) under identical conditions showed a similar early decline in permeate flux (Figure 2B). In contrast, after IgA/IgM depletion by Fractogel TMAE chromatography, sequential Planova 35N→20N nanofiltration processed 60 mL within 140 min under the same operating pressure, with stable filtrate accumulation and no marked decline in flow. The corresponding average permeate flux was 25.7 L·m⁻²·h⁻¹ (Figure 2C, Table 2). When Planova S20N (0.001 m²) was used after 35N prefiltration, the steady permeate flow increased to ∼1.5 mL·min⁻¹ compared with ∼0.5 mL·min⁻¹ for Planova 20N. This corresponds to over three-fold increase in permeate flux (∼90 vs 25.7 L·m⁻²·h⁻¹) (Figure 2D, Table 2), with stable flow and no evidence of progressive membrane fouling over the recorded volume range.

**Figure 2.**
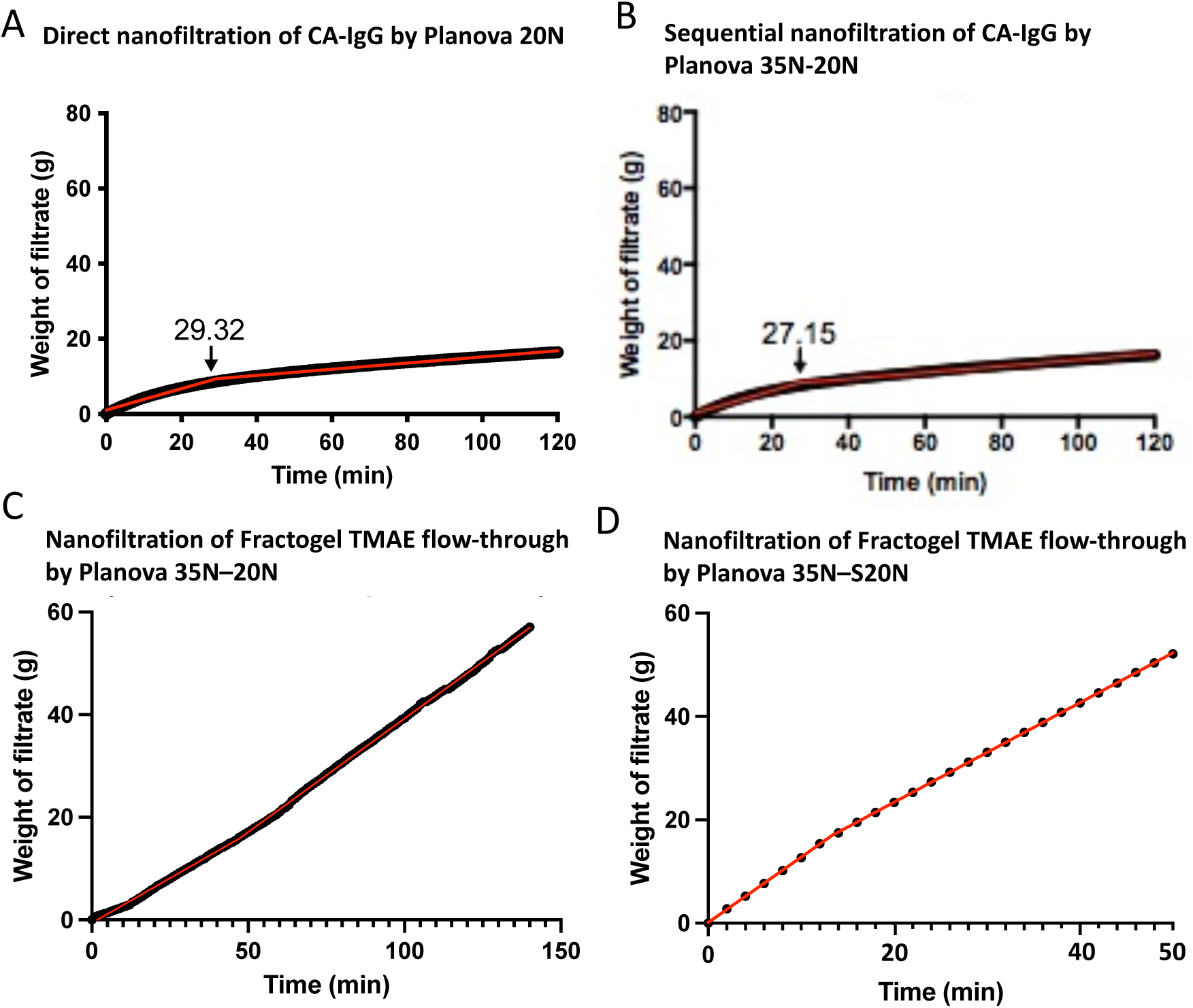
Effect of IgA/IgM depletion on Planova nanofiltration performance of caprylic acid–purified IgG (CA-IgG). (A) Direct nanofiltration of CA-IgG using Planova 20N (membrane area 0.001 m²) shows an early decline in permeate flux after ∼29 min, yielding only ∼16 mL of filtrate after 120 min (0.8 kgf/cm²). (B) Sequential filtration of CA-IgG using Planova 35N followed by 20N (0.001 m² each) under identical conditions does not prevent early flux decline (after ∼26–27 min) and results in a similar filtrate volume (∼15–16 mL after 120 min). (C) After Fractogel TMAE chromatography (IgA/IgM-depleted flow-through), sequential Planova 35N→20N filtration shows stable filtrate accumulation, indicating improved filterability after upstream removal of fouling components. (D,E) Volume–time plots used to derive permeate flow rates for the Fractogel TMAE flow-through during sequential filtration using Planova 35N→20N (D) and Planova 35N→S20N (E). Under the same feed conditions, the S20N run shows a higher steady permeate flow (∼1.5 mL/min) than the 20N run (∼0.5 mL/min). In panels A–C, cumulative filtrate mass (g) is plotted versus time; overlaid traces represent repeated runs.

**Table 2.** Effect of IgA/IgM depletion on nanofiltration performance of caprylic acid–purified IgG (CA-IgG) using Planova membranes. Filtration conditions and outcomes are shown as filtrate volume, filtration time, membrane area, and calculated average permeate flux. Average permeate flux (L·m⁻²·h⁻¹) was calculated as filtrate volume / (filtration time x membrane area). For the Planova S20N run, permeate flux was calculated from the stable permeate flow rate (≈1.5 mL. min^-1^) and the membrane area. “-” indicates no IgA/IgM depletion prior to filtration, and “√” indicates that IgA/IgM were removed by Fractogel TMAE chromatography before nanofiltration.

| **Filtration step** | **IgA/IgM removal** | **Filtrate volume (mL)** | **Filtration time (min)** | **Membrane area ($\text{m}^2$)** | **Average permeate flux ($\text{L}\cdot\text{m}^{-2}\cdot\text{h}^{-1}$)** |
| --- | --- | --- | --- | --- | --- |
| Planova 20N | - | 16 | 120 | 0.001 | ~8 |
| Planova 35N → 20N | - | 15–16 | 120 | 0.001 | ~8 |
| Planova 35N → 20N | √ | 60 | 140 | 0.001 | 25.7 |
| Planova 35N → S20N | √ | 60 | 40 | 0.001 | ~90 (flow $\approx 1.5 \text{ mL}\cdot\text{min}^{-1}$) |

*Average permeate flux was calculated as the total filtrate volume divided by the product of filtration time and membrane surface area. For Planova S20N, permeate flux was calculated from a stable permeate flow rate of approximately $1.5 \text{ mL}\cdot\text{min}^{-1}$*

### IgG Integrity and Particle Size

DLS analysis of the IgG preparations (Figure 3A-B) showed Z-average hydrodynamic diameters of 11–12 nm and PdI values <0.2 after chromatography and/or nanofiltration, indicating a narrow distribution of scattering species and no evidence of increased aggregation under the tested conditions. In Fractogel TMAE flow-through fraction (Figure 3A), two DLS peaks were observed, with a higher PdI (0.397) and a larger Z-average for the main peak (15.95 nm). Following sequential 35N and 20N nanofiltration, the minor secondary DLS peak at larger hydrodynamic diameters, was no longer detected indicating removal of a population of large particles or IgG aggregates. After Planova 35N-20N nanofiltration, the PdI decreased to 0.142 and the Z-average decreased to 11.04 nm, consistent with improved sample homogeneity. A similar Z-average (≈12 nm) indicated a monomeric IgG was maintained after S20N filtration (Figure 3B). NTA showed that sequential nanofiltration with both Planova 20N or S20N reduced the concentration of detectable subvisible, light-scattering particles/aggregates in the IgG preparations (Figure 3C). All measurements were performed within a comparable tracking density (∼30–33) particles per frame. The particle counts decreased from 2.84 × 10¹¹ particles/mL (35N) to ∼2.6–2.7 × 10¹⁰ particles/mL (35N–20N or 35N–S20N) (≈11-fold reduction), yielding low particle counts after filtration. Together, DLS and NTA indicate that both sequential Planova nanofiltration reduces low-abundance large particles or aggregates while maintaining a dominant IgG-sized population.

**Figure 3.**
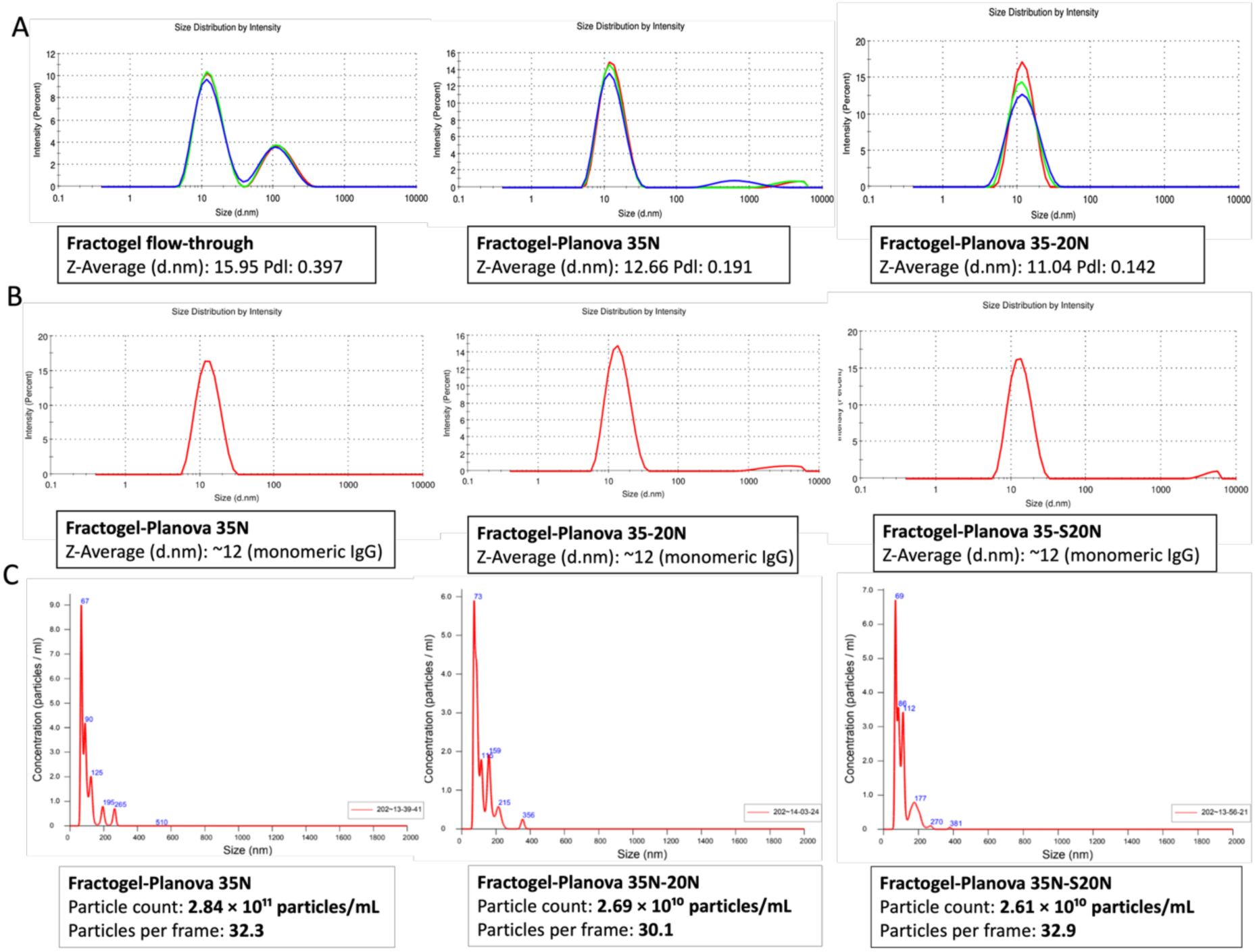
DLS and NTA profiles of the apparent size distribution and subvisible particulate content of IgG intermediates after Fractogel TMAE chromatography and Planova nanofiltration. (A) Dynamic light scattering (DLS) intensity-weighted hydrodynamic size distributions of the Fractogel flow-through and of products after Planova 35N filtration and sequential Planova 35N→20N filtration. Z-average hydrodynamic diameter and polydispersity index (PDI) are shown below each plot. (B) Representative DLS intensity distributions for the corresponding products (Fractogel–35N, Fractogel–35N–20N, and Fractogel–35N–S20N). (C) Nanoparticle tracking analysis (NTA) of the same IgG solutions showing the concentration and apparent size distribution of subvisible, light-scattering particles (IgG aggregates/particulates); total particle concentration (particles/mL) and measurements at comparable tracking density or particles per frame (∼30–33) are indicated in each panel. The NTA data show a marked reduction in detectable subvisible particles after Planova 20N or S20N filtration.

SDS–PAGE (Figure 4) supported preservation of the IgG profile after sequential Planova nanofiltration and indicated depletion of higher-molecular-weight species. Under non-reducing conditions, all samples displayed a predominant band at ∼150 kDa consistent with intact IgG, with no detectable low-molecular-weight degradation products and no additional prominent high-molecular-weight bands.

**Figure 4.**
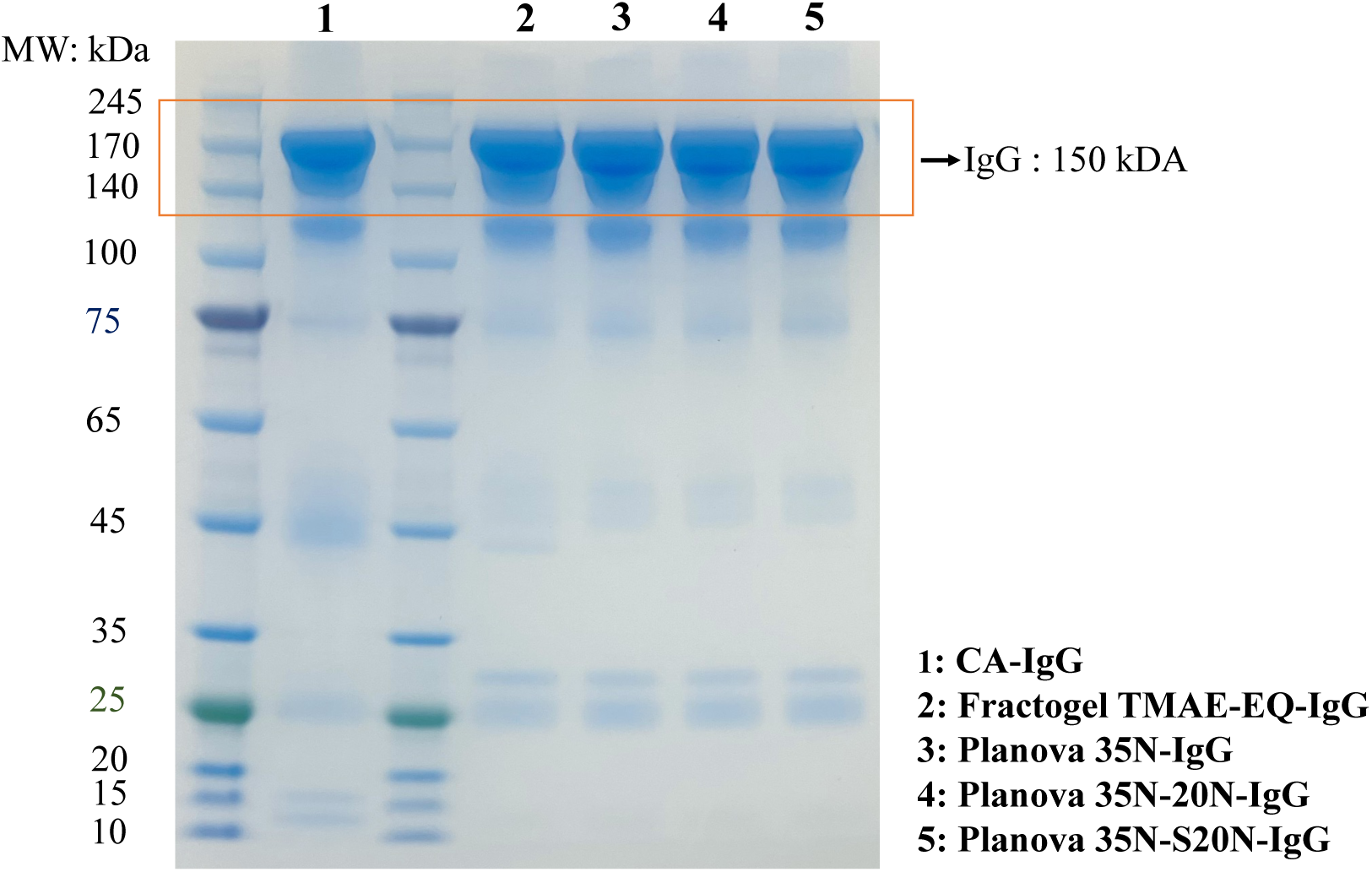
SDS–PAGE profile of IgG intermediates under non-reducing (conditions before and after IgA/IgM depletion and Planova nanofiltration. All samples show a dominant band at ∼150 kDa consistent with intact IgG and depletion of higher-molecular-weight species after sequential Planova nanofiltration. Lane assignments: (1) caprylic acid–purified IgG (CA-IgG); (2) Fractogel TMAE (M) flow-through (EQ-IgG); (3) Planova 35N filtrate; (4) sequential Planova 35N→20N filtrate; (5) sequential Planova 35N→S20N filtrate. Molecular weight markers (kDa) are shown on the left.

## Discussion

This study shows that adding an anion-exchange chromatography step to selectively remove IgA and IgM enables efficient nanofiltration of CA-purified IgG using small-pore virus removal filters. In our previous work, Fractogel® TMAE anion-exchange chromatography also provided high IgG recovery (mean 96.9%), increased IgG purity from about 75% in the CA intermediate to >96%, and removed >96% of IgA and IgM. IgG subclass distribution was preserved, and column performance remained consistent over 200 cycles, supporting robustness and scalability (Cheng et al., 2021). That previous study also addressed the risk of procoagulant contaminants, in particular activated factor XI (FXIa), which has been implicated in thromboembolic events associated with some IVIG products (Food and Administration, 2010; Germishuizen et al., 2014; Roemisch et al., 2011). FXI/FXIa was not detectable after CA treatment, and no thrombin generation activity was observed in the purified IgG fraction in thrombin generation assays (Cheng et al., 2021). In addition, the downstream anion-exchange step provided an extra safeguard by removing residual procoagulant activity, consistent with the absence of thrombin generation in the final IgG preparation (Cheng et al., 2021). Together, these results support the CA–Fractogel® TMAE process as both scalable and aligned with current expectations for thrombosis-related quality attributes.

From a virus safety perspective, combining CA treatment of CPP with nanofiltration using 20 nm-class filters (e.g., Planova 20N or S20N) provides two complementary virus reduction steps. CA treatment inactivates lipid-enveloped viruses (El-Ekiaby et al., 2015), while 19 nm nanofiltration has been widely validated for removal of both enveloped and non-enveloped viruses (Roth et al., 2020b). Reported log-reduction factors depend on process conditions and model viruses, but CA treatment can provide strong inactivation of enveloped viruses under appropriate pH and concentration conditions (Dichtelmuller et al., 2002; Radosevich and Burnouf, 2010), and 19 nm nanofiltration can provide robust removal of small non-enveloped viruses in validation studies (Roth et al., 2020b). Our results add a practical point: selective IgA/IgM depletion before nanofiltration is critical to maintain filter performance and to obtain an IgG preparation with improved homogeneity that should be suitable for therapeutic development.

CA precipitation at pH around 5.5 has long been used to manufacture intact equine antivenom immunoglobulins, mainly by precipitating non-immunoglobulin proteins while keeping immunoglobulins in solution (Rojas et al., 1994). Similar approaches have also been applied to hyperimmune or convalescent plasma-derived immunoglobulins (e.g., SARS-CoV-2) (Ali et al., 2021a). CA inactivates enveloped viruses through disruption of lipid membranes, and this activity is influenced by pH because the nonionized form is more lipophilic and partitions more readily into viral envelopes (Dichtelmuller et al., 2002; Korneyeva et al., 2002; Li, 2019). Importantly, while CA enriches IgG and contributes to enveloped virus inactivation, it does not substantially deplete IgA and IgM (Ali et al., 2021a; El-Ekiaby et al., 2015). The persistence of these higher-molecular-weight immunoglobulins, particularly IgM, can strongly impair the capacity and throughput of small-pore nanofilters. In our experiments, Planova 20N showed rapid flux decline when IgA and IgM were present, whereas filter capacity dramatically increased after IgA/IgM removal by Fractogel® TMAE chromatography.

Chromatography and nanofiltration also have practical advantages because both can be implemented at different scales (Burnouf, 1995; Roth et al., 2020b). Columns and disposable nanofilters can support small batches in blood establishments and can be scaled up for larger pools in centralized facilities. This flexibility could support routine production of polyvalent IgG and may also be applicable to hyperimmune IgG products (e.g., anti-Rh or anti-HBs). The same platform could also be used for convalescent IgG production during outbreaks when rapid, local manufacturing is needed and plasma volumes are limited. Product quality is further supported by the absence of increased aggregation and the improved homogeneity indicated by DLS, together with reduced subvisible particulate content measured by NTA. Removal of IgA may also reduce the likelihood of adverse reactions in immunodeficient patients with anti-IgA antibodies (Rachid and Bonilla, 2012).

From an implementation perspective, the basic CA precipitation process is accessible to blood establishments as already demonstrated by the preparation of clinical-grade products in Egypt (Elalfy et al., 2017; Selim et al., 2025), but successful deployment of chromatography and nanofiltration (especially when not performed in closed systems) requires GMP-aligned upgrades, including controlled water and buffer preparation, validated column cleaning/sanitization, environmental controls, and well-defined in-process monitoring. These needs can be addressed through modular equipment and disposable, pre-packed columns to allow phased implementation in LMICs. Optional additional stepwise steps such as anti-A/anti-B reduction or final concentration for subcutaneous formulations can be added depending on local resources, regulatory requirements, and clinical needs (Cheng et al., 2021).

This study is limited by the scale of testing and the need for broader process validation and clinical evaluation. Nonetheless, the results provide a practical roadmap for local IgG production in LMICs to reduce plasma wastage and improve access to essential therapies for immunodeficiencies and outbreak response. This platform aligns with the World Health Organization’s stepwise approach to expanding access to plasma-derived medicinal products in LMICs (WHO, 2021). Finally, quality-assured plasma remains a prerequisite: donor selection, testing, traceability, and cold-chain management must meet established quality requirements for safe and consistent immunoglobulin manufacture (Weinstein, 2018; WHO, 2005). Public–private partnerships can facilitate technology transfer, training, and sustainable scale-up.

In summary, combining CA precipitation with scalable anion-exchange chromatography and small-pore nanofiltration provides an option to produce high-quality IgG at small-to-medium scale, including in resource-constrained settings. This strategy could help LMICs build fractionation capacity and expand access to essential PDMPs within a WHO-recommended stepwise framework (WHO, 2021).

## Conclusion

This study shows that CA precipitation of plasma followed by Fractogel TMAE anion-exchange chromatography and 19 nm-class nanofiltration can generate a high-purity IgG preparation from CPP with two complementary virus-reduction steps. The critical enabling factor was IgA/IgM depletion prior to nanofiltration, which restored filterability and yielded an IgG solution with improved homogeneity and low levels of detectable subvisible particulates. This modular process can be implemented at different scales and provides a practical option for local production of polyvalent or specialty IgG, including in settings where industrial fractionation is not available, provided GMP-aligned controls can be established. Technology transfer and training, potentially through public–private partnerships, will be important for sustainable implementation.

## Acknowledgments

The authors thank Asahi Kasei Medical for financial support.

## Data availability

The data that support the findings of this study are available from the corresponding author upon reasonable request. Data will be made available on request.

